# The role of myosin contractility as a molecular determinant of traction force regulation

**DOI:** 10.1101/2022.08.30.505821

**Authors:** Debsuvra Ghosh, Subhadip Ghosh, Abhishek Chaudhuri

## Abstract

Cells generate traction forces on the extracellular matrix to crawl forward. A complex molecular machinery is involved in the generation, transmission, and transduction of cellular forces inside and outside of cells. The molecular clutch hypothesis, with motors as rudimentary force generators, has been beneficial in modelling the distinctive biomechanical roles played by the components of this machinery. In this paper, we propose an analytical model that incorporates the active dynamics of myosin motors and establishes their roles in regulating the traction force in an experimentally accessible parameter space. As the parameters pertaining to molecular determinants are varied, we show that the system traverses between diverse states of stabilities - from decaying oscillations to self-sustaining limit cycles. The hallmarks of motor-clutch models like load-and-fail dynamics and shift in traction optima are successfully encapsulated. Modulating myosin activity in our model via different pathways exhibits striking shifts in optimal stiffness, providing excellent agreement with experiments and additional testable predictions.

## 1 Introduction

The bulk of cellular functionalities depends on the consistent exertion of biomechanical forces and their subsequent channelling through intracellular molecular machinery towards extracellular environments. The key differentiator in carrying out such processes is the stiffness of the extracellular matrix (ECM) – it directs cell proliferation [1], regulates the differentiations of cells [2], controls migrations [3] and moulds cell mortality [4], to name a few. Apart from substrate stiffness, biological ECMs including fibrin [5], collagen and tissues [6] exhibit viscoelastic properties via molecular mechanisms like protein unfolding, release of entangled states and breaking of fragil crosslinks [7] – leading to stress relaxation effects. Migrations guided by rigidity, i.e., *Durotaxis* require cells to probe the ECM to establish directional motion and regulate them further. The molecular machinery inside the cell responsible for such movements comprises a force-generating actomyosin complex, adaptor proteins that carry these forces, and transmembrane heterodimers like integrins which establish linkages to the ECM [8, 9, 10, 11, 12]. Together they form the cellmatrix adhesion complex and manifest a meticulous multilaminar protein assembly [13] controlling specific cellular adhesion to the ECM. Myosin-mediated contractility, coupled with polymerisation of actin pressing against the cellular membrane, triggers a persistent flow of F-actin called *retrograde flow* [14]. Experimental evidence suggests that retrograde flows are inversely proportional to the speed of cell migration [15], indicating that the force transferred to the ECM counters myosin contractility, resulting in a reduced retrograde flow. The dynamic nature of the engagement and release of various components of this actin cytoskeleton-ECM complex led to the coining of the term *molecular clutch* [16] that transmits forces to ECM/substrate and mechanosenses the response to turn them into regulatory signalling pathways.

Active sliding of the actin filaments by the myosin motors is one of the most crucial force generation mechanisms within the cell-matrix adhesion complex that controls force fluctuations in clutches [17], modulations of mechanosensing, and cell migrations [18, 19, 20]. Extensive in-vivo and in-vitro experimental studies provide important insights into the workings of individual myosin motors [21] and their collective performance through the actomyosin complex [22]. Myosin activity is pivotal in focal adhesion initiation [23, 24], maturation [25] and speed of migration [26]. Experiments using traction force microscopy have demonstrated that myosin inhibition increases traction force transmission for a specific range of stiffness [27]. Theoretical models [28, 29, 30, 31, 32, 33, 34, 35, 36, 37, 38, 39, 40] have explored the interplay between the different dynamic elements of the cell migration machinery. Biophysical modelling based on the molecular clutch hypothesis led to the fundamental prediction of a *biphasic force–substrate stiffness relationship* [32] with a traction peak at an *optimum stiffness* [34]. Further, dual parameter changes in the model allowed for a broad range of the stiffness optima, which is shown to be sensitive to the expression of molecular motors and clutches [33, 37]. However, motors are relegated to static force generators with a stalling mechanism in all such models. The effect of myosin attachment/detachment to/from the actin cytoskeleton and their transient engagement with the focal adhesion machinery is expected to significantly affect traction force dynamics.

In this study, the distinct role of myosin II activity in the clutch-substrate sector is incorporated into a theoretical model [17] of traction force dynamics on a visco-elastic ECM. The activity of myosin II motors proteins (MPs) is expressed both in terms of their turnover dynamics with F-actin and through the velocities of the attached motors. We set forth a time-dependent coupled ordinary differential equation representation of the model with the goal of understanding the time evolutions of operative units of the adhesion complex. An extensive analysis of the stabilities of actomyosin-clutch-substrate sectors leads to a collection of dynamical phases governing the system behaviour. We show that the model correctly predicts the biphasic stiffness dependence consistent with experiments and earlier theoretical models. Modulation of motor activity in (i) the number of molecular motors, (ii) motor velocity, and (iii) motor attachment/detachment dynamics leads to dramatic shifts in the stiffness optima. While some of the results show excellent agreement with experimental results, others are predictive and leave scope for further experimental and theoretical studies.

This article is organised in the following manner. We introduce the theoretical model in section 2 by stating the dynamical equations. In section 3, we analyse the time evolution equations using linear stability analysis and enumerate different dynamical regimes by means of various phase diagrams in terms of active and passive parameters. We specially focus in the linearly unstable regime, where a bi-phasic behaviour is observed as the substrate stiffness is changed. Finally, in section 4 we conclude by presenting current findings and future possibilities.

## 2 Model

We posit a 1-dimensional arrangement of F-actins in the proximity of force-generating myosin II motors, force-transmitting molecular clutches, and a compliant substrate. One end of the motors is rigidly anchored, and they apply forces to the F-actin bundle by attaching themselves via the opposite ends, inducing a *retrograde flow.* Similarly, the clutches can attach to the F-actin using a reversible mechanism on one edge while permanently attached to the substrate through the other. The accumulation of mechanical force on the attached clutches ushers in a *traction force* that is balanced by the deformation of the pliant substrate as shown in Fig. 1(b).

**Figure 1:**
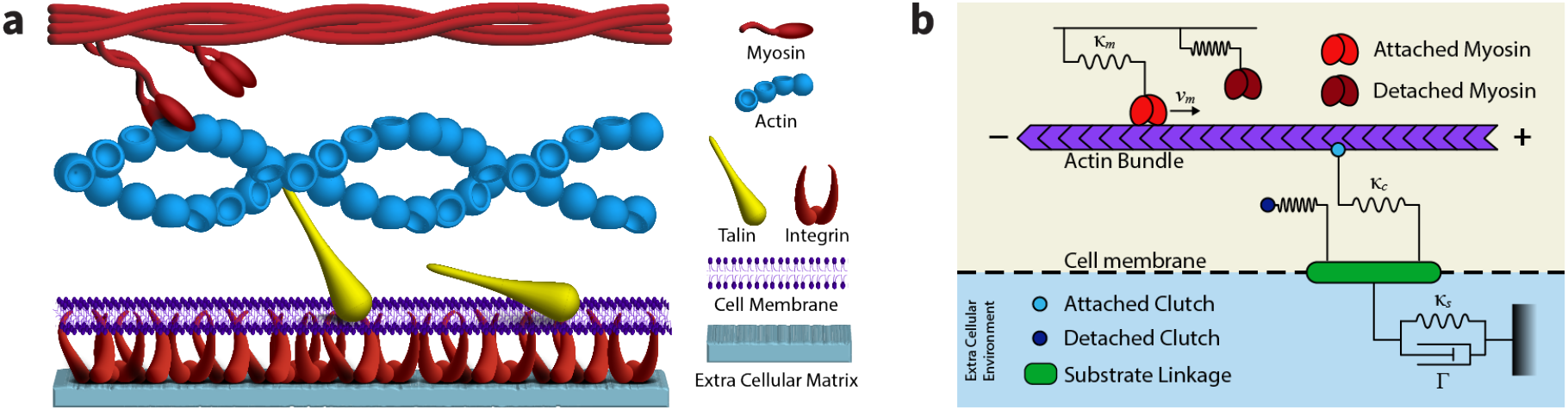
(a) A cartoon depicting serially connected cell migration machinery including myosin motors and actin filament which constitutes the cross-linked cytoskeletal network. Force exerted via this network is transmitted to the extracellular matrix/substrate through focal adhesion (FA), using adaptor proteins (talin) and transmembrane proteins (integrin). (b) A schematic of a motor-clutch model considering the myosin motors and clutches as elastic springs. The rigid actin filament with ± points to anterograde/retrograde directions. The substrate is considered to be a viscoelastic Kelvin-Voigt material. The figures were adapted from Ghosh *et al.* [17] and modified to fit the current context.

Following the treatment from our previous study [17], we follow a coarse-grained approach where MPs and clutches are considered as Hookean springs with load-dependent attachment-detachment dynamics, i.e. their numbers at connected state (*n_m_* & *n_c_*) are determined by the tension acting upon them 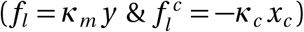.

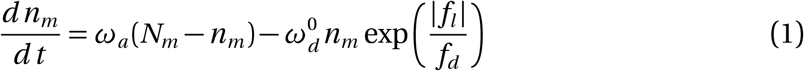

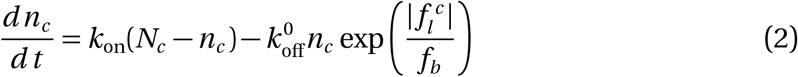

MPs attach with rate *ω_a_* and has a bare detachment rate 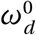, while clutches connect with rate *k*_on_ and possess a bare detachment rate 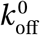. The mean extensions of MPs and clutches are *y* and *x_c_* respectively. The tensile forces, *f_l_* and 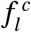, coupled with the viscous forces that the clutches experience, are then stabilised through a *force-balance* relation,

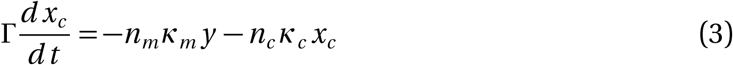

where the viscous clutch force is designated by viscous coefficient *Γ*.

The clutch and the substrate sectors are treated as separable blocks as they are connected in series. We model the substrate as a visco-elastic material according to Kelvin-Voigt description as a viscous dashpot connected to a Hookean spring in parallel (Fig. 1). Correspondingly, a new force-balance equation concerning the substrate arises,

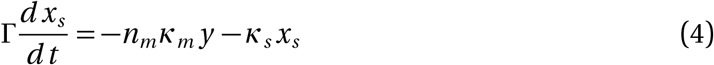

where *x_s_* is the substrate extension, and *κ_s_* is the stiffness. The F-actin retrograde flow gets equalised by the motor-clutch-substrate sector deformation dynamics,

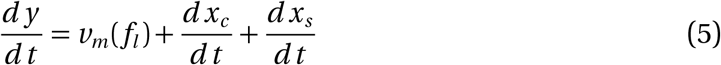

Connected MPs move along the actin filament with a velocity *v_m_*(*f_l_*) that depends on the load force experienced by the F-actin, expressed with a piecewise linear force-velocity relation,

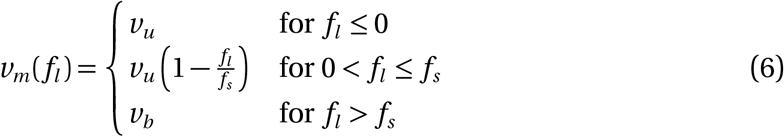

Hence, we arrive at a closed set of differential equations describing the dynamics of our system with Eqs. (1–5).

We proceed with a comprehensive analysis of the stability of the MP–F-Actin–clutch–substrate system, focusing on the variations of substrate mechanical properties. Table 1 describes the biophysical parameters employed in our model and provides their experimental values. The dynamical system, comprising of Eq. (1), (3), (4), (5) and (2), is converted to dimensionless form with appropriate time, length, velocity and force scales as 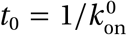, *l*_0_ = (*k_b_T/k*_on_Γ)^1/2^, *v*_0_ = *l*_0_*k*_on_, and *f* = (*k*_on_Γ*k_b_T*)^1/2^, respectively and consequently using *τ* = *t*/*t*_0_, 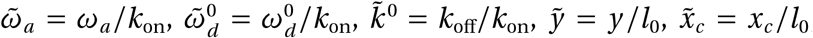, 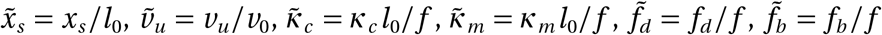 and 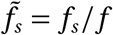 (see Appendix A). Motor association/dissociation dynamics is varied using a turnover ratio defined by 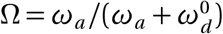.

**Table 1:** Physical parameters present in the system

| Parameter | Symbol | Values |
| --- | --- | --- |
| Motor attachment rate | $\omega_a$ | $40 \text{ s}^{-1}$ [41] |
| Motor detachment rate | $\omega_d^0$ | $350 \text{ s}^{-1}$ [41] |
| Total number of motors | $N_m$ | 100 |
| Clutch attachment rate | $k_{\text{on}}$ | $1 \text{ s}^{-1}$ [32] |
| Clutch detachment rate | $k_{\text{off}}$ | $0.1 \text{ s}^{-1}$ [32] |
| Back velocity | $v_b$ | $-0.2256 \mu\text{m s}^{-1}$ [41] |
| Stall force | $f_s$ | $4.96 \text{ pN}$ [41] |
| Detachment force | $f_d$ | $2.4 \text{ pN}$ [42] |
| Clutch bond rupture force | $F_b$ | $6.25 \text{ pN}$ [33] |
| Motor spring constant | $\kappa_m$ | $0.3 \text{ pN/nm}$ [41] |
| Clutch spring constant | $\kappa_c$ | $0.03144 \text{ pN/nm}$ [43] |
| Viscous friction coefficient | $\Gamma$ | $893 k_B \text{ Ts}/\mu\text{m}^2$ [44] |

## 3 Results

To characterise the linear stability of the scaled differential equations, a set of steady-state solutions or fixed points is determined as follows, 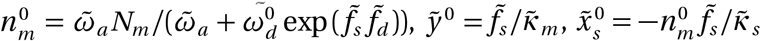, and 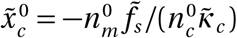. Accordingly, the steady-state extension of the connected clutches is calculated by solving the following transcendental equation,

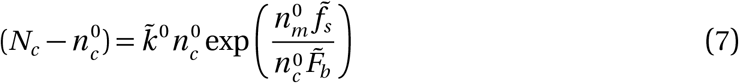

Linear stability analysis is utilised to study the evolution of small perturbations away from the fixed points. To that end, our system can be linearised to obtain a matrix-operator form, 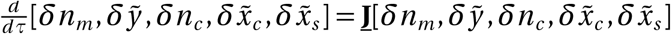, where 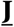 is a 5×5 Jacobian matrix (see Appendix B) whose eigenvalues dictate the linear stability of our system. These eigenvalues are determined from the solutions of a fifth-order characteristic polynomial equation,

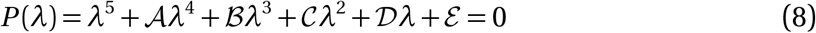

whose coefficients 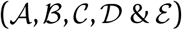 are calculated using the aforementioned scaled parameters (see Appendix B).

Inspecting the nature of this polynomial equation alone, without explicit numerical solutions, provides us with a trove of information about the properties of its roots, and therefore their stabilities. It is noted that 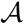 is always real-positive (+ℝ) while the remaining coefficients allow for changes in their signs with parametric variations. The biophysical nature of the parameters ensures that they are real-valued and thus each of the coefficients is a real number as well. This puts a constraint on the five roots of the quintic polynomial *P*(*λ*) that one of them must be real-valued. Considering the roots as *λ_k_* (*k* = 1…5), denoting the eigenvalues, the requisite real root is labeled *λ*_1_. The remaining four roots can have the following combination: (1) all real-negative (–+) eigenvalues indicating stable nodes with exponentially decaying perturbations; (2) two –ℝ and two +R that correspond to unstable nodes where perturbations grow exponentially; (3) two –ℝ and two complex conjugate eigenvalues with negative real part i.e. –*α*±*iβ* that points to stable spiral phases with damped oscillations and finally (4) two +ℝ and two complex conjugate eigenvalues with positive real part i.e. *α* ± *iβ* corresponding to unstable spiral phases with growing oscillations. With the aforementioned restriction in place, we proceed to determine closed form expressions for the viable phase boundaries present in the system.

The saddle-node bifurcation arising in Eq. (7) leads to two branches – one stable and another unstable, which can be characterised by checking the signs of the derivative equation around the bifurcation point (Figure 6). The presence of this bifurcation on the number of connected clutches points to initial condition-dependent dynamics, i.e. a system starting above (below) the unstable branch will eventually lead to stability (instability) in the system. We will discuss the situations in detail to understand how they lead to diverse dynamical phases.

Confirming 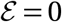 as a phase boundary, and the existence of an unstable branch at 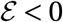, we focus on the stable branch 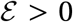. A collection of phase boundaries emerges in this branch with varying signs of 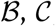, and 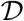. Mathematical derivations of their closed form expressions are given in Appendix C. A phase diagram is then presented on the *N_c_* – *v_u_* plane (Figure 2) to encapsulate the varied stabilities in the system.

**Figure 2:**
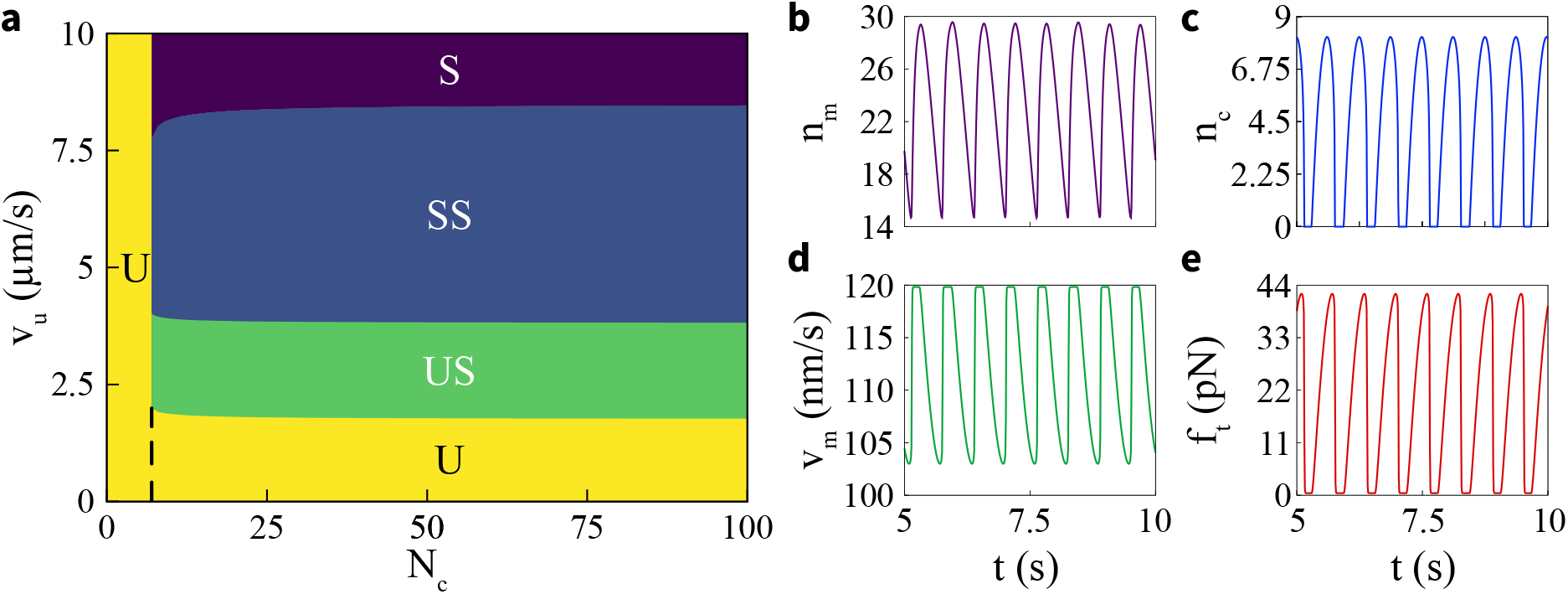
Phase diagram on *N_c_–v_u_* plane: The dynamical phases predicted by stability analysis are illustrated using coloured regions with labels - 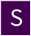 stable, 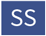 stable spirals, 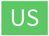 unstable spirals, and 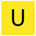 unstable. The unstable region is divided into two parts - one originating at very low *N_c_* on the left of the dashed line owing to absence of fixed points and another due to 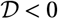. **Slip-stick behaviour on the unstable branch:** MP-clutch-substrate system initiated below the 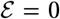 boundary leads to a cycle of progressive loading and cascading failure of the clutch sector, as shown in (c). Fluctuations in *n_m_* (b) lead to rhythmic stalling and free-flowing of F-actin retrograde velocity (d), which in turn give rise to an oscillating traction force *f_t_* (e).

### Load-and-fail dynamics

The model dynamics begin with MPs actively connecting to polar actin filaments. As both MPs and clutches begin to bind with their respective attachment rates, a force transduction pathway is engaged and myosin contractility begins to tug on the substrate. As a result, a compliant substrate becomes deformed and the total applied force is distributed among the connected clutches. The system, when initiated below 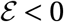 near the unstable branch in Figure 6, starts with a severely low number of connected clutches. Due to relatively higher attachment rates and ready availability of MPs, they rapidly bind and build up a tremendous amount of force that is to be distributed among very few connected clutches. This leads to a cascading failure of clutches and the force exerted on the ECM via clutches immediately goes to zero. With no connected clutches present, the clutch attachment mechanics begins to dominate and the cycle continues. The actomyosin-clutch sector here is balanced in such a manner that the traction force rhythmically loads before eventually failing. This behaviour is known as *load-and-fail* or *stick-slip* [32]. It becomes evident from the clutch and traction force dynamics in Figure. 2, as they fluctuate between zero to positive contributions.

The stable branch, at 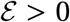, leads to a richer array of dynamics. As the initially connected number of clutches is safely larger than the relevant fixed point at the bifurcation point, they are not likely to be influenced by the unstable branch. At a constant substrate stiffness, higher *v_u_* leads to a prompt, large extension of MPs and thus stalling of their motions. The retrograde flow comes to a minimum and the clutch–substrate sector attains a force balance, resulting in a stable (S) state. An increase in *N_c_* leads to a strengthened clutch-substrate sector that forces MPs to stall at relatively higher *v_u_*. At lower *v_u_*, the MP sector is sufficiently weakened to allow for a *load-and-fail* cycle between the MP and clutch sectors[17]. The presence of a compliant substrate allows for these cycles to stabilise over a larger parameter space (Figure 2).

### Constraints on motors recover basic motor-clutch model behaviour

To validate our model, we begin with a constrained case where myosin motors do not have attachment-detachment dynamics and are not stretchable. In this scenario, they are merely force generators with active velocities. The dynamical equations are modified to adapt to these changes resulting in a system of 3 differential-algebraic equations. All motors present in the system are always connected and the total force exerted by them is balanced separately by the aggregate force from the connected clutches and by substrate deformation,

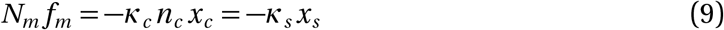

where *f_m_* is the average force applied by each motor.

Myosin velocities are still attenuated by the restoring force from the substrate via a force-velocity relation which is now independent of motor extension,

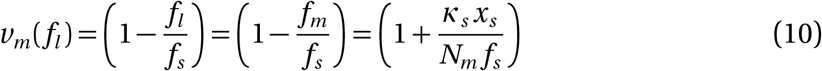

In the absence of a motor extension, the load-dependent active velocity of myosin is matched by the rate of average deformation of connected clutches and the substrate,

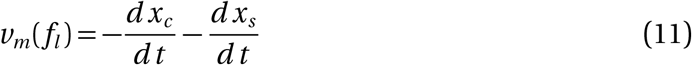

These equations coupled with the clutch number dynamics from Eq. (2) give us the constrained *Differential-Algebraic equations* governing the system. We proceed with numerically solving the equations to study the nature of the traction force generated in the system. In line with the previous studies [32, 33, 34], numerical solutions to this constrained system points to an *optimal substrate stiffness*, at which the average traction force is maximum. Conversely, the mean retrograde flow of the actin filament i.e. *v_m_*(*f_l_*) is at its minimum. Increasing the total number of motors and clutches together leads to a significant shift in this optimal stiffness as evident from Fig. 3.

**Figure 3:**
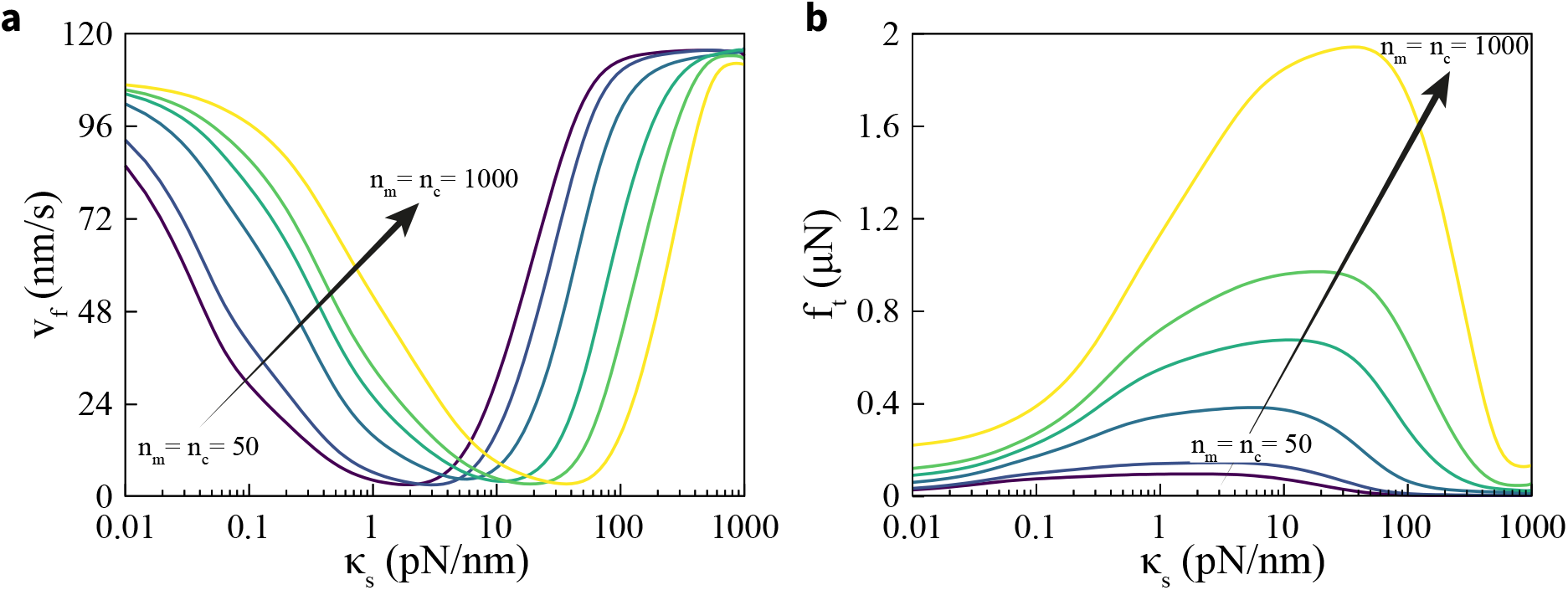
Shifting of optimal stiffness: A joint increase in the number of motors and clutches shifts the optimal stiffness towards higher values. As *n_m_* = *n_c_* is increased from 50 to 1000, (a) the minimum mean retrograde flow is shifted from around 2 pN to 40 pN. (b) records the same shift in maximum mean traction force which increases substantially with large numbers of motors and clutches.

### Regulation of MP-clutch sector explicates force transmission on ECM

Apart from the simultaneous motor-clutch sector contributions through tuning of their population, several other parameters affect the mechanosensitivity of the clutch-substrate sector. Analysing modulations of the *loading rate* serves as an excellent indicator, which is the speed of forces building up in the engaged clutch ensembles. Being the force generators in the system, MP parameters directly influence the loading rates. Inhibiting myosin contractility leads to a lowering of loading rate which should shift the zone of optimal stiffness towards higher substrate stiffness. However, experiments using mouse embryonic fibroblasts suggest that inhibition of MPs using various concentrations of myosin inhibitor blebbistatin leads to a rise in force transfer to ECM in certain rigidity ranges, resulting in shifting of traction force peak to higher rigidities with gradually decreasing amplitudes [27]. Our model captures this behaviour as we lower the total number of MPs present in the system gradually (Fig. 4 (a)). It is noted that beyond a threshold of rigidity *κ_s_*, a declining *N_m_* leads to transfer of optimal force to increasing substrate rigidity.

**Figure 4:**
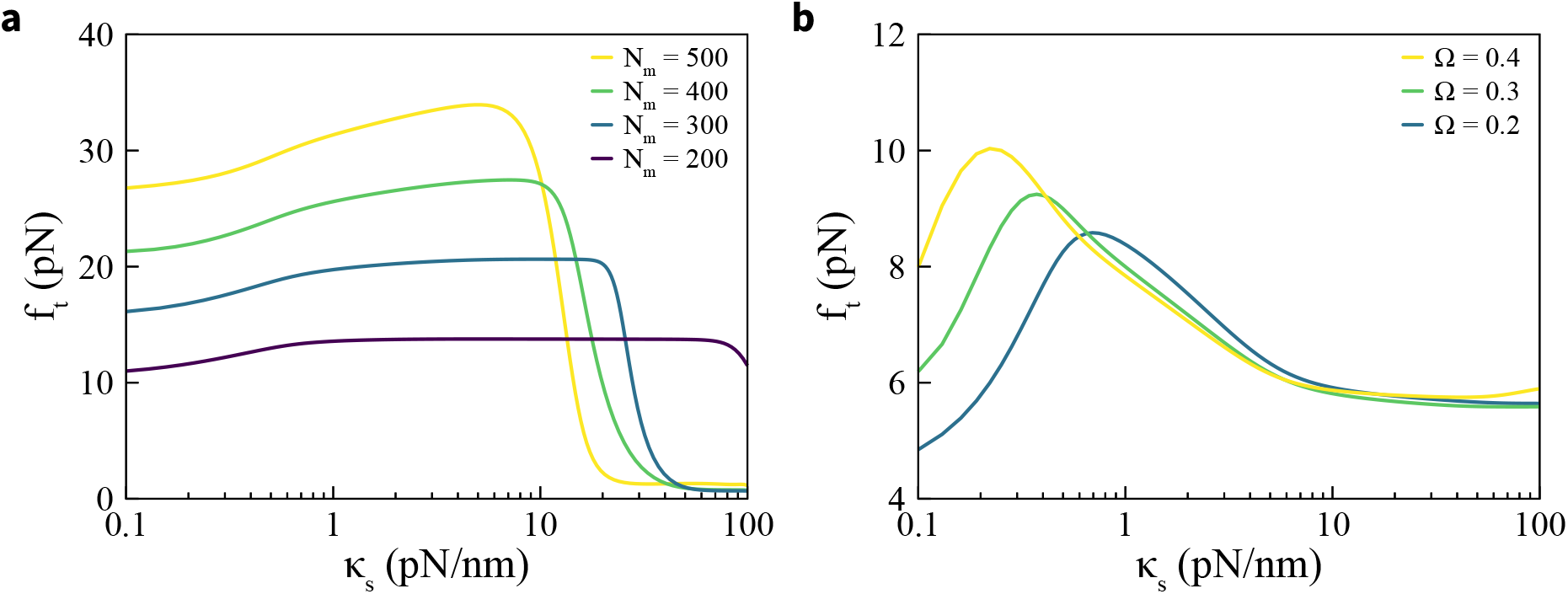
Myosin regulates optimal stiffness: Tuning myosin motor parameters involved in mechanosensitivity leads to dramatic shifts in optimal stiffness. (a) Inhibition of myosin activity moves the traction peaks towards higher substrate stiffness, increasing transmission of traction forces beyond a rigidity threshold of about 1 pN/nm. (b) Increasing motor attachment rate *ω_a_* through the motor duty ratio *Ω* at *v_u_* = 1 *μ*m/s results in progressively lower stiffness optima, strengthening the amplitudes of traction force transmission.

Motor turnover dynamics is another hallmark of myosin activity. Increase in both motor active velocities and their attachment rates can enhance the self-sustaining oscillations in focal adhesions over a larger parametric space [17], while variations in motor detachment rates can lead to a range of frequencies for the tugging forces as observed in experimental setups [18, 45]. Our model predicts a sharp rise in traction forces with increasing *ω_a_*, with the maxima settling towards higher substrate stiffness, as seen in Fig. 4 (b). Stronger motor attachment rates allow substantial portions of *N_m_* to end up in connected state, thereby elevating the loading rate which in turn transfers the traction maxima towards lower rigidities.

Clutch sector activity can be hindered by an increase of clutch dissociation or by pruning the total number of clutches available to the system. Employing inhibitory agents like GPen-GRGDSPCA peptide (GPen) can effect such changes on integrin binding mechanism [27] or different types of integrins having a range of binding/unbinding rates can be used [46]. On the other hand, altering the number of available clutches *N_c_* is also possible [47]. From a computational perspective, our model allows for changes in the related parameters such as *N_c_* and *k*_off_ to capture the behavioural shift in traction forces. With increasing *k*_off_ and the dwindling number of connected clutches *n_c_*, the overall force exerted by the motors are distributed among fewer clutches and therefore the traction peak veers to a lower *κ_s_*. This is visualised in Fig. 5 (a) where gradual increase in *k*_off_ from 0.1 Hz to 0.75 Hz hauls the optimum towards 1 pN/Nm of rigidity. Furthermore, a decrease in *N_c_* from 80 to 20 also facilitates similar shift as evident from Fig. 5 (b).

**Figure 5:**
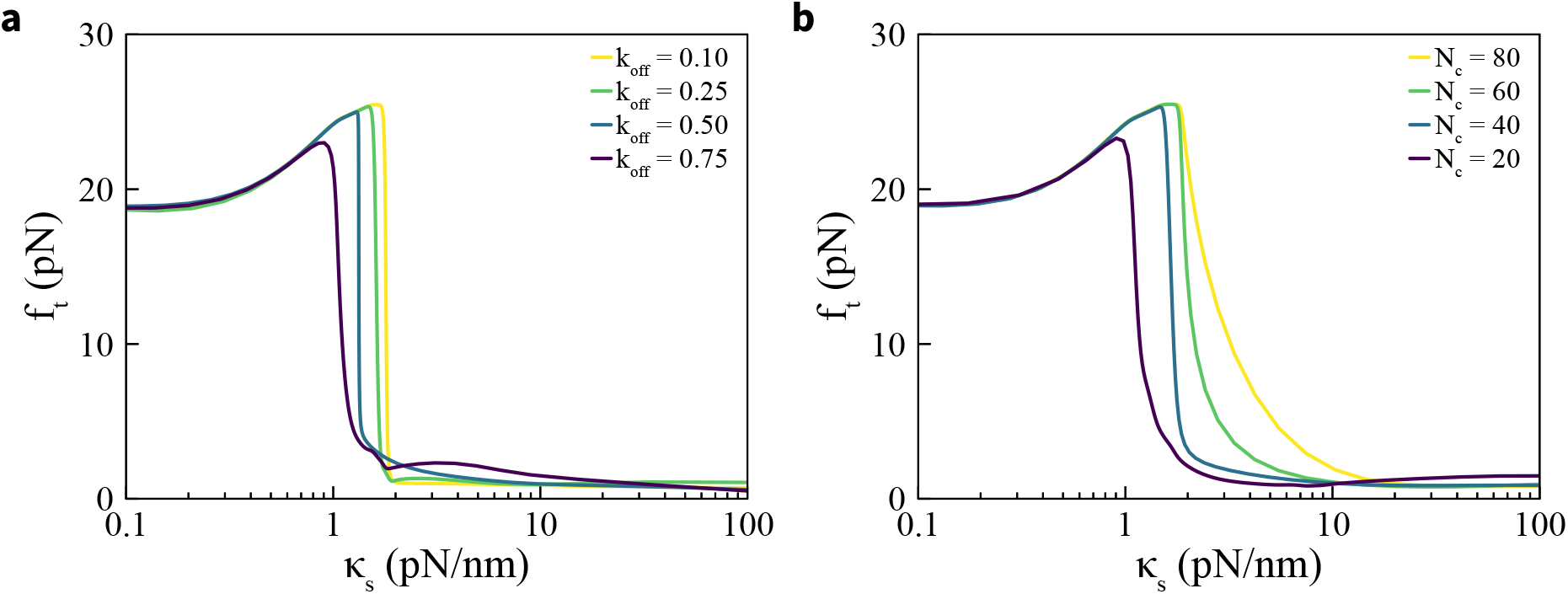
Stiffness optima depends on clutch parameters: (a) Increasing clutch detachment rate *k*_off_ results in progressively lower stiffness optima, veered towards lower substrate stiffness. (b) Increments in the total clutch number *N_c_* strengthen the clutch sector and ensues enhanced optimal stiffness moving towards higher rigidities.

**Figure 6:**
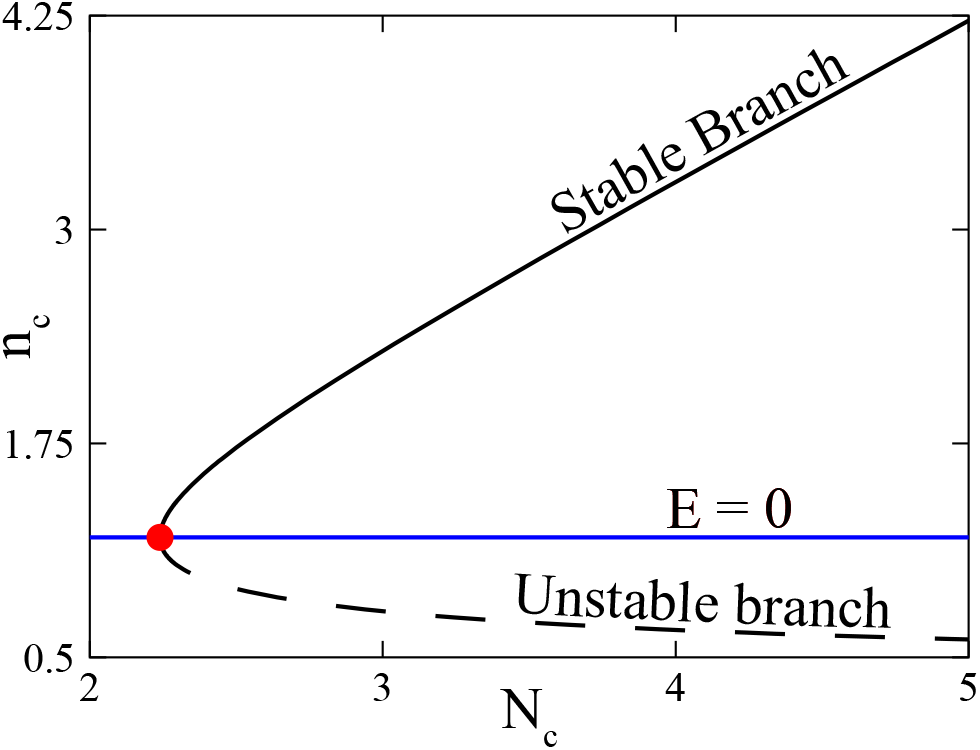
Bifurcation Diagram: Fixed point equation of *n_c_* in Eq. (7) is transcendental, which causes a *saddle-node bifurcation* with two distinct branches – solid black line indicating the stable branch and dashed curve signifying the unstable one. They part from the bifurcation point marked by a red dot.

## 4 Discussion

Optimal stiffness is a recurrent phenomenon across cell types in diverse environments. Preconditioning of the vascular smooth muscles via mechanical cues can produce notable changes in gene expressions, leading to possible enhanced mechanical strength. Variations in ECM stiffness provide the smooth muscle cells with such cues and control the speed of their random motilities in a biphasic manner, indicating an optimal region [48]. Excessive accumulation of ECM proteins causes liver fibrosis, which can advance to liver cirrhosis and eventual failure. Transient elastography is a non-invasive method to gauge fibrosis via liver stiffness measurement (LSM), and optimal stiffness cutoff values provide a metric for fibrosis progression [49, 50]. Pathological instances aside, optimal rigidity also provides anatomical evolutionary advantages. Lobsters evolved to have pore canals that have regular honeycomb-type geometry with thick walls and maximum possible stiffness for the same porosity fraction. They ensure optimal cuticle stiffness and maximal fluid transport, enabling swift mineralisation without losing structural integrity [51]. Examples like these not only point to the significance of a stiffness optimum in multitudes of biological systems, but they also stipulate a wide variability in the domains of optimality. Therefore, understanding the role of relevant and experimentally accessible parameters in tuning the optimum is paramount.

MP and clutch sector contributions in force generation and transduction are well documented. In particular, an in-vivo experiment conducted on U251 glioma cells by Bangasser *et al.* [37] showed simultaneous changes in the total number of MPs and clutches alter the optimum stiffness for a cell. In their setup, polyacrylamide hydrogel (PAG) substrates with a range of Young’s modulus and surface-bound collagen type I were used to monitor the migration of human U251 glioma cells. As myosin II motors in the cytoskeleton were partially inhibited using blebbistatin, cyclo(RGDfV) was used to inhibit the integrin–substrate adhesion partially. This drug treatment combination shifted the actin flow minimum towards lower ECM stiffness.

While experimental results signify their roles, there is a dearth of biophysical models that integrate these components along with their individual dynamics. Earlier theoretical attempts, while successful in predicting rigidity optima, did not account for the variations in the forces originating from the stress fibres [32, 34] as they failed to consider the role of dynamic attachment/detachment to/from the actin cytoskeleton on myosin contractility and therefore on the force regulation machinery in such systems [33, 52]. A number of stochastic models of the traction force generation introduced minor modifications to [34] and fitting of parameters around experimental data [46, 27]. Our model substantiates that an extensive set of dynamical phases are present in the system for a wide biologically feasible range of parameter values. Further joint adjustments to the MP/clutch numbers shift the stiffness optima towards higher rigidity (Fig. 3) as shown by Bangasser *et al.* [37].

The key reason to focus on the MP sector emanates from the fundamental factor that force loading rate governs the response of the molecular clutches, and MPs play a crucial role as force generators in cells. In this work, we provide an analytical framework that recognises the importance of MP activity and association-dissociation mechanics while enabling us to examine the traction force fluctuations and modulations of filament velocities. We acknowledge the significance of internal (myosin contractility, clutch types and their numbers) and external (viscoelastic nature of substrate) factors influencing the loading rate to present a modular, extensible reaction network that is sufficiently simple yet captures the intricacies introduced by these factors.

Durotaxis is guided by the transmission and transduction of cell-ECM forces; therefore, it is an optimistic assumption that biophysical models which predict the nature of these forces would also be successful in characterising cell migration. However, traction forces produced by cells are manifolds larger than migratory requirements. In experimental setups, a migrating cell produces tractions that nullify themselves when summed up within the margins of environmental noise, concluding that they cannot be considered propulsive [53]. Nevertheless, cell protrusions are directly dependent on actin polymerisation rate and retrograde flow. Following our observation in Fig. 3, traction is maximised at optimal stiffness while minimising retrograde flow simultaneously. Therefore, cells with minimal retrograde flow should be the quickest to migrate for a specific actin polymerisation rate Multicellular or collective durotaxis works akin to single cell migration, i.e. they prefer to advance towards stiffer substrate rigidity. Characterisation of intercellular junctions would lead to an obvious extension of our model with cell-specific reaction networks tugging at the ECM while maintaining an overall force balance throughout the interconnect. Once the inherent stochasticity is considered, it will be interesting to explore emergent phenomena like *collective intelligence* that indicates grouped systems performing more efficiently than their components [54, 55].

## Acknowledgements

We acknowledge the use of the computing facility at IISER Mohali. S.G. acknowledges QuantiXLie Centre of Excellence, a project co-financed by the Croatian Government and European Union through the European Regional Development Fund - the Competitiveness and Cohesion Operational Programme (Grant No. KK.01.1.1.01.0004).

## Appendix A: Dimensionless Equations

Considering the physical parameter values described in Table 1, we continue to convert our dynamical equations dimensionless as prescribed in the main text. The characteristic scales for length, time, velocity, and force are calculated as *t*_0_ = 1 s, *l*_0_ = 33 nm, *v*_0_ = 33 nm/s, and *f* = 0.125 pN.

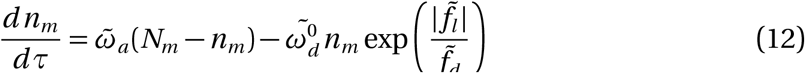

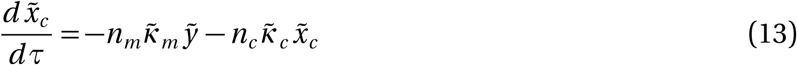

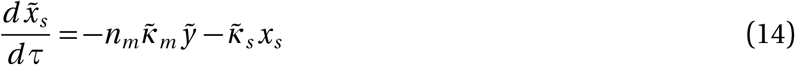

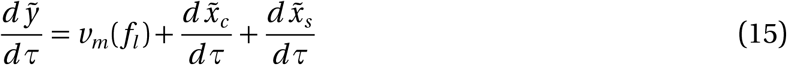

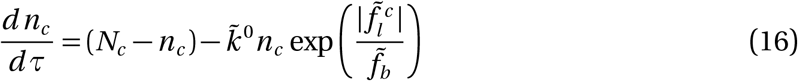

## Appendix B: Jacobian

A Jacobian matrix 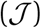 is determined to calculate the linearisation about the fixed points of the system mentioned in main text. There are 5 dynamical variables and corresponding differential equations, and therefore the Jacobian matrix is of the order 5 × 5. It contains 25 elements as shown in Eq. (17)

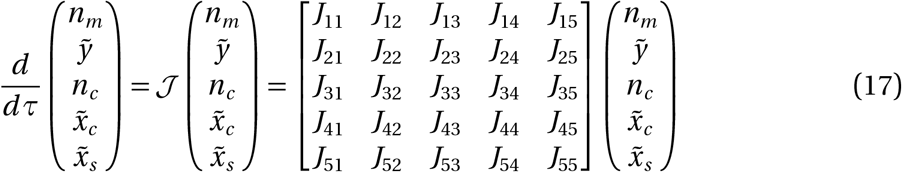

The elements of the Jacobian matrix, *J_ij_*, are explicitly computed and the complete matrix is shown below,

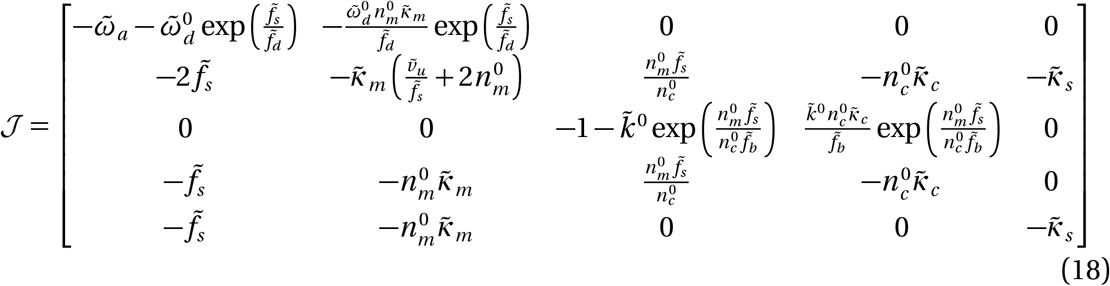

The characteristic polynomial, as mentioned in the Eq. (8), is now a fifth-order polynomial equation where 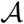 is the negative trace of 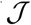 i.e. 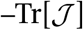 and 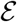 is the determinant or 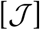. A complete list of coefficients are given below,

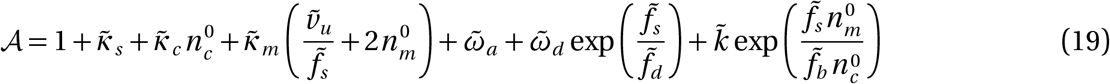

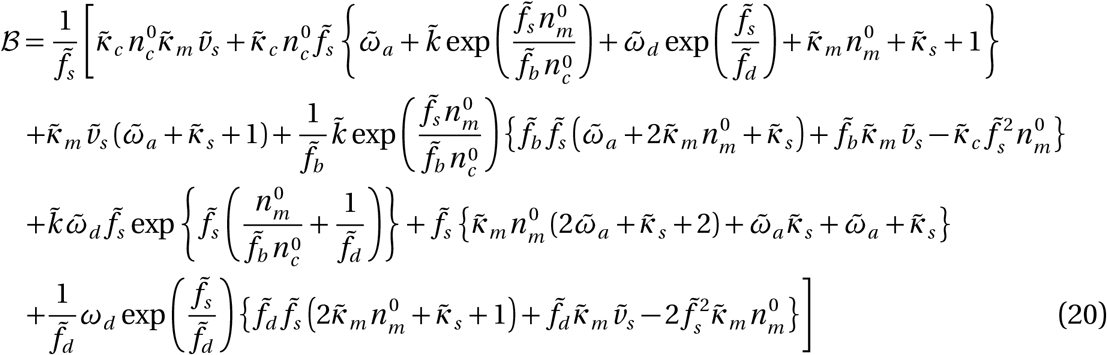

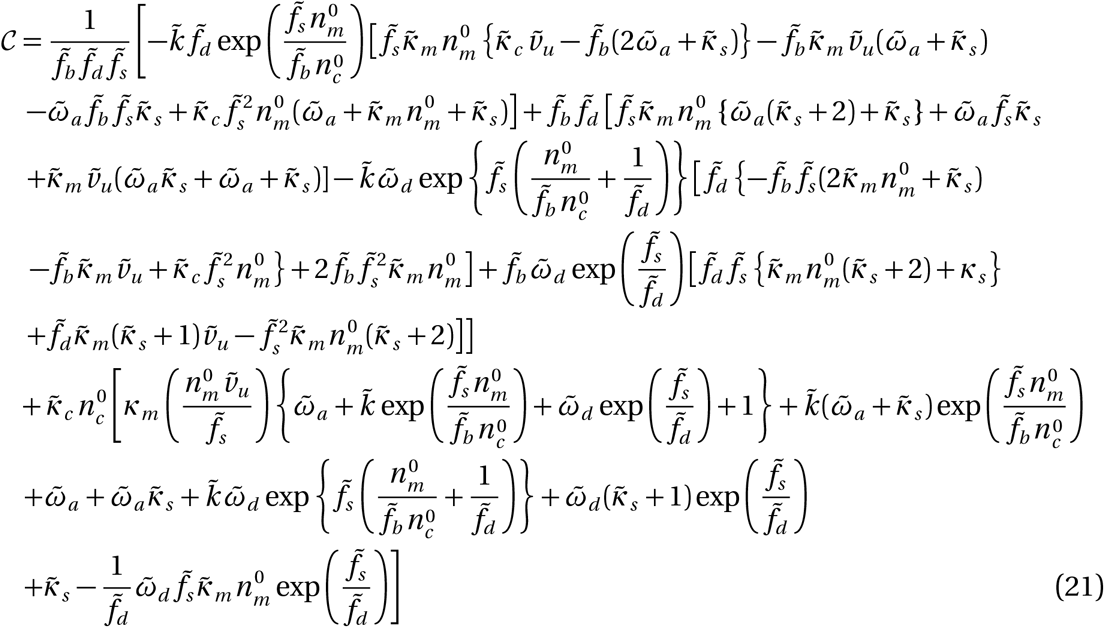

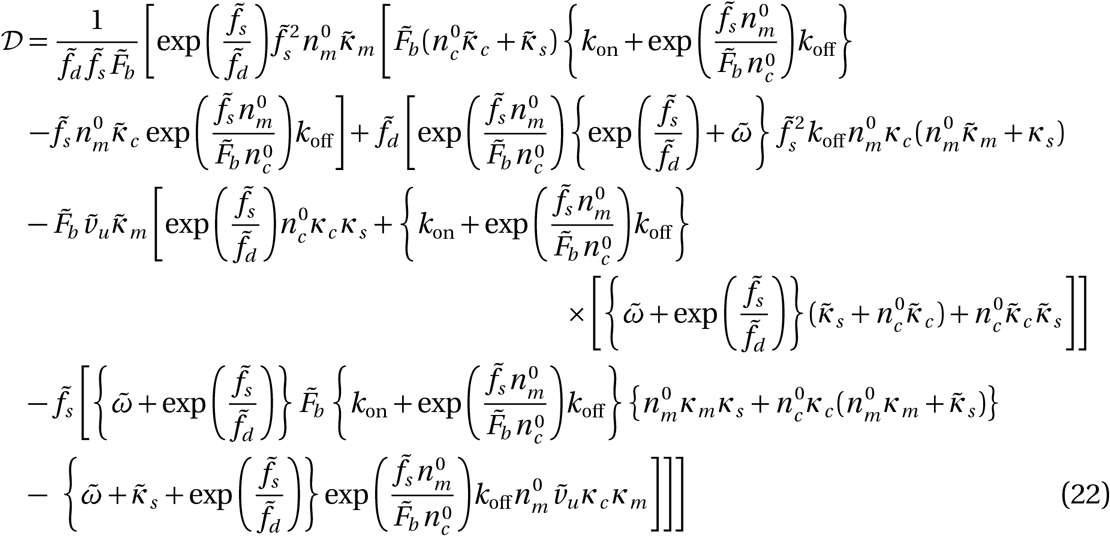

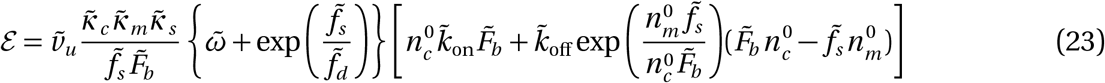

Nature and properties of the eigenvalues are dependent on the sign of the coefficients 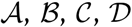, and 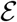. We explore the algebra of polynomial equations to ascertain the features of the roots that they possess, which, in turn, provides us with the dynamical phases without explicitly solving the differential equations governing the system. In the following section, we shall detail a method to systematically determine the characteristics of algebraic roots of a real-valued polynomial equation.

## Appendix C: Phase boundaries

### 4.1 Saddle-node bifurcation

The transcendental Eq. (7) produces a saddle-node bifurcation containing two branches. Their stabilities can be characterised by checking the sign of their derivatives along the branches. The location of a *bifurcation point* can be computed by solving Eq. (7) while setting its derivative with respect to 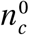 to zero simultaneously. This derivative equation is as follows,

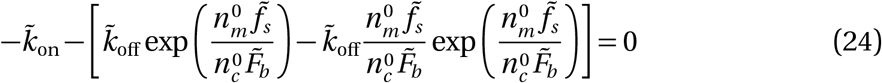

### 4.2 Phase boundary between stable and unstable branches delineated by saddle-node bifurcation

It is trivial to show that Eq. (24) is the same as a phase boundary equation at 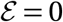,

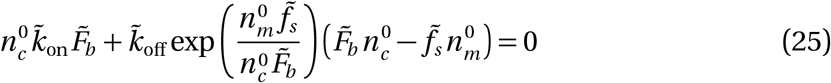

A simplified form of Eq. (25) can be obtained using Eq. (7) as follows,

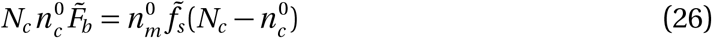

### 4.3 Phase boundary between stable and unstable spirals

The Hopf bifurcation boundary is given by the following expression,

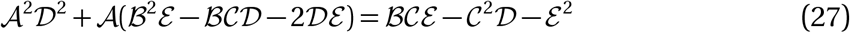

### 4.4 Phase boundary between (un)stable and (un)stable spirals

This boundary has the following expression,

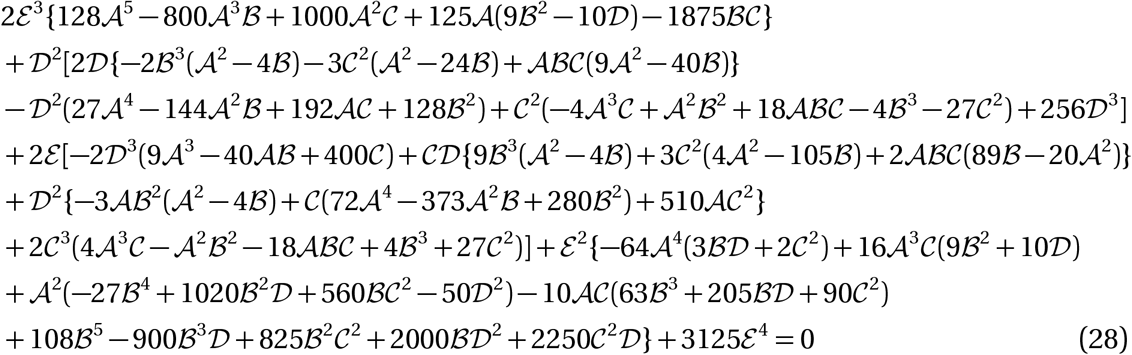

